## Supplementary figures and images for "Redistribution of fragmented mitochondria ensure symmetric organelle partitioning and faithful chromosome segregation in mitotic mouse zygotes"

### Actin expression during preimplantation development.tif

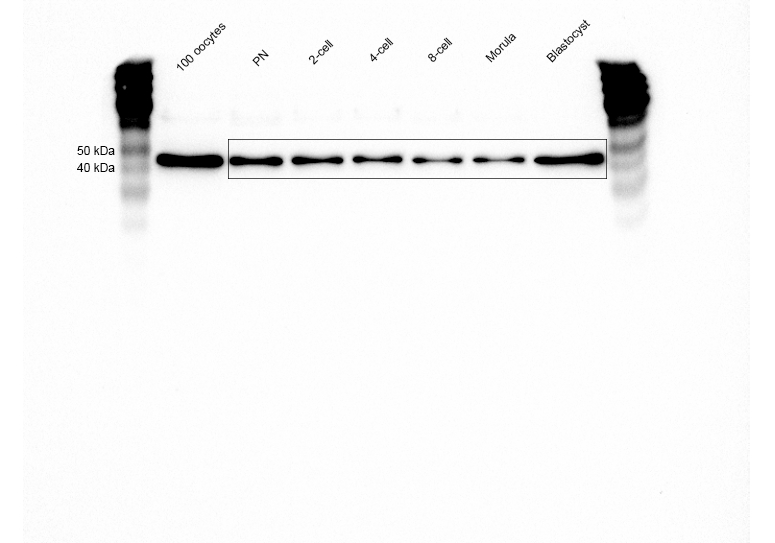

### Actin expression during preimplantation development_raw.jpg

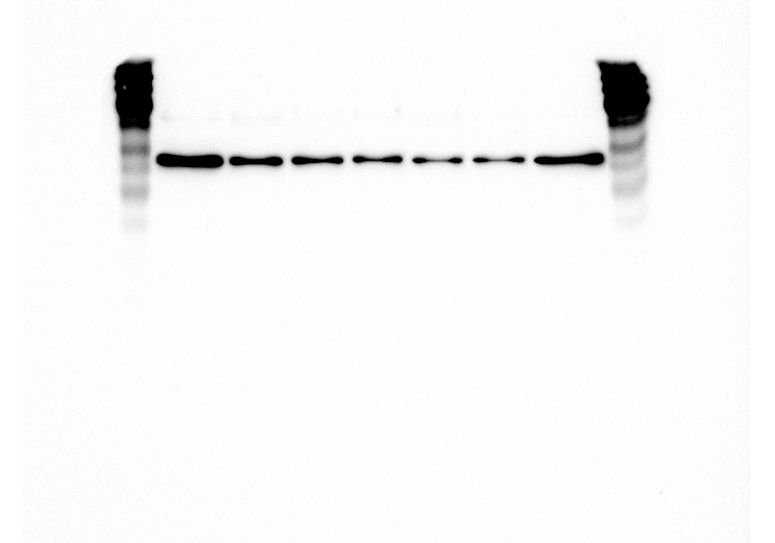

### Actin expression rescue raw.tif

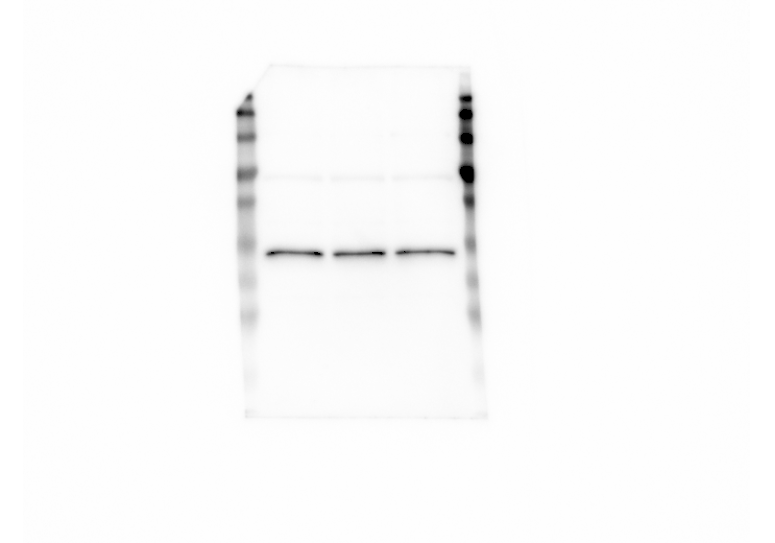

### Actin expression rescue.tif

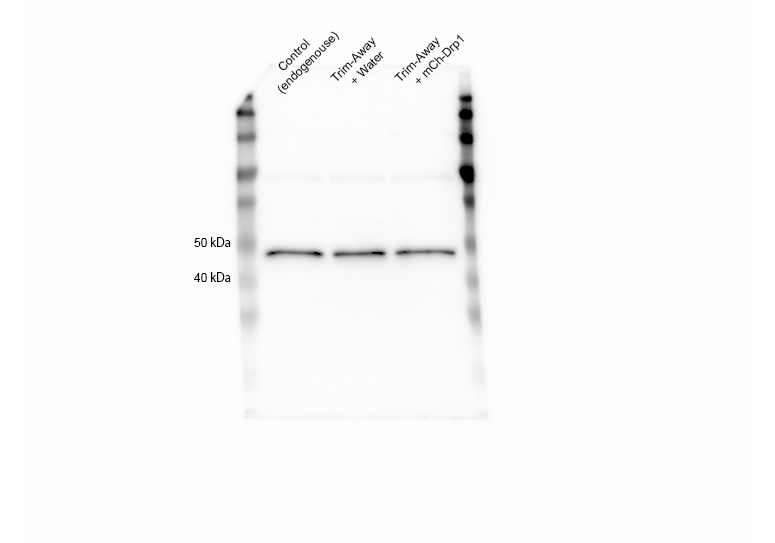

### Actin expression Trim Away raw.tif

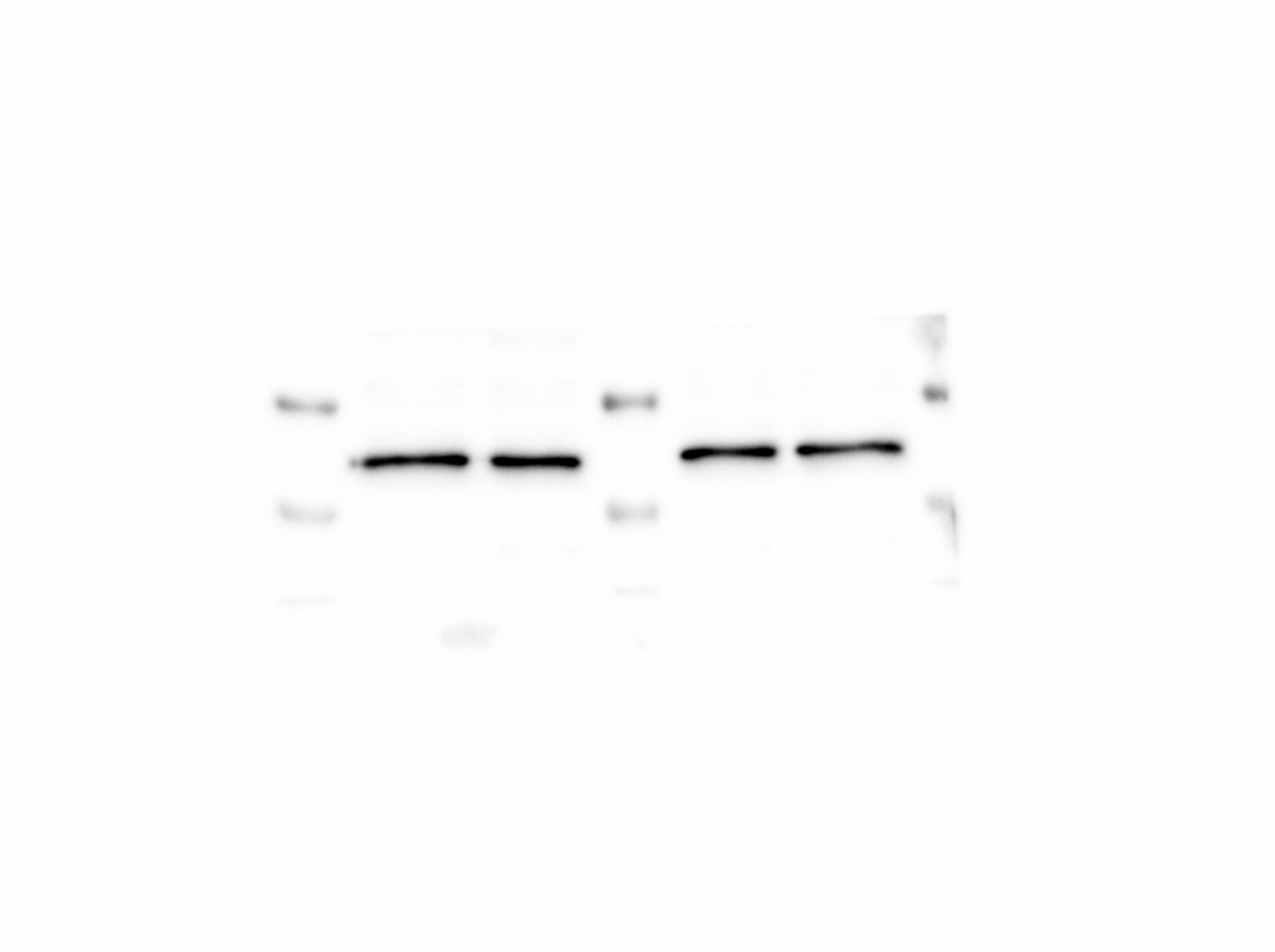

### Actin expression Trim Away.tif

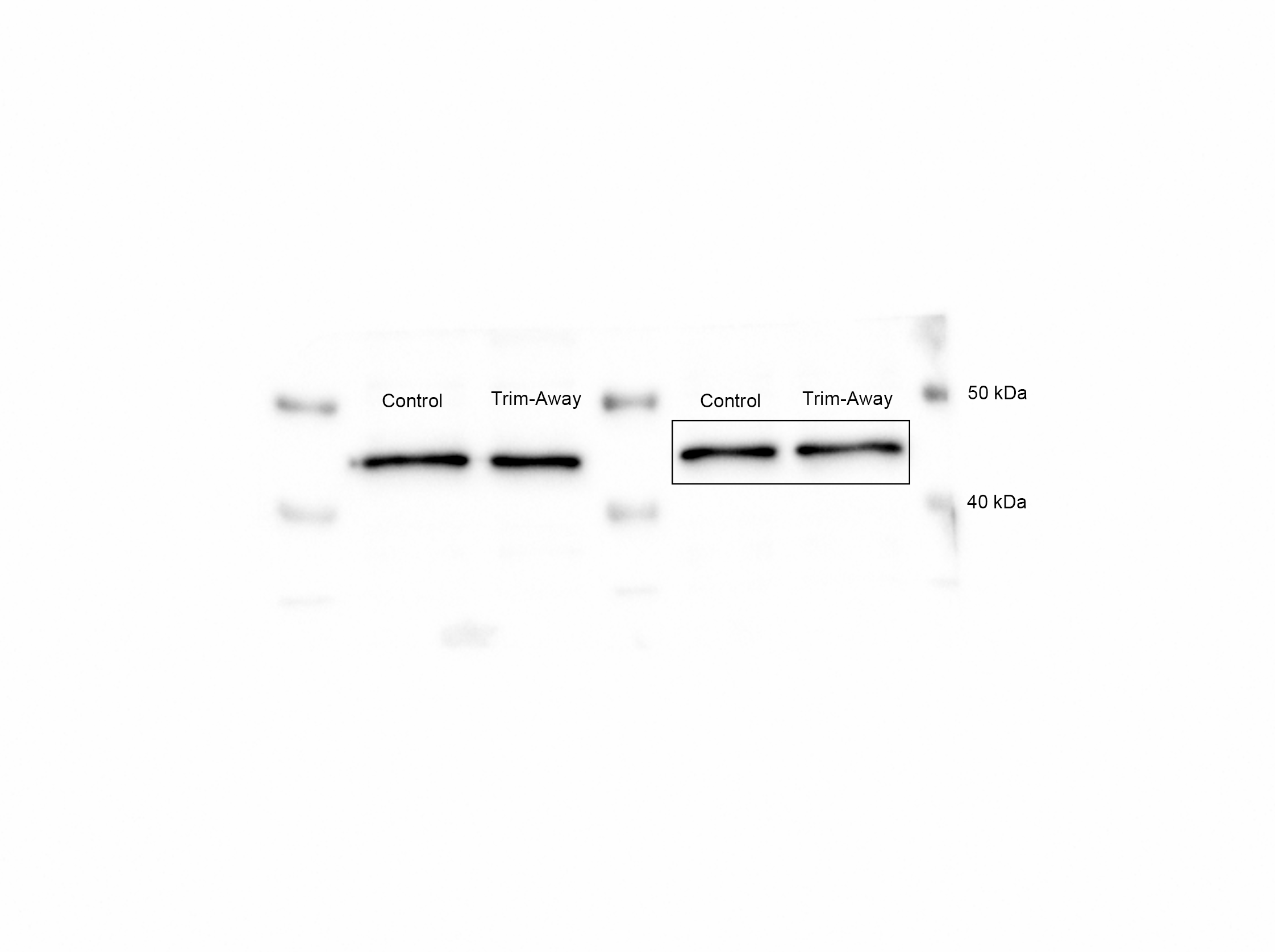

### Actin expression Trim Away.tif

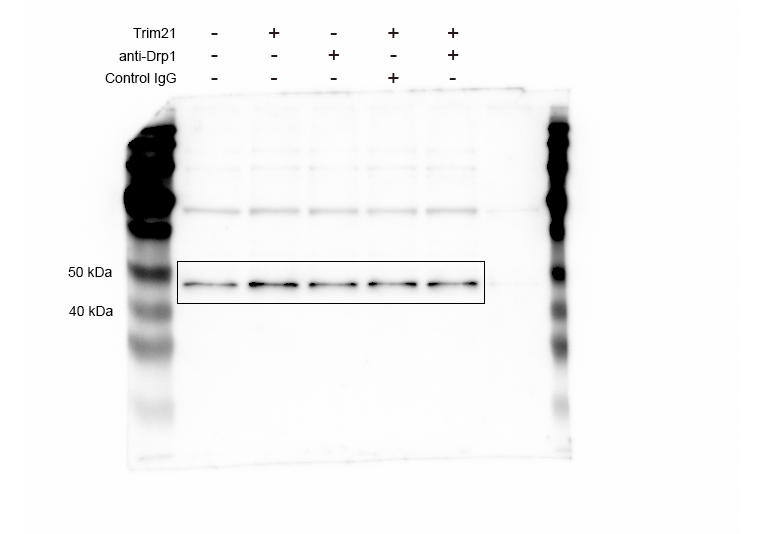

### Actin expression Trim Away_raw.tif

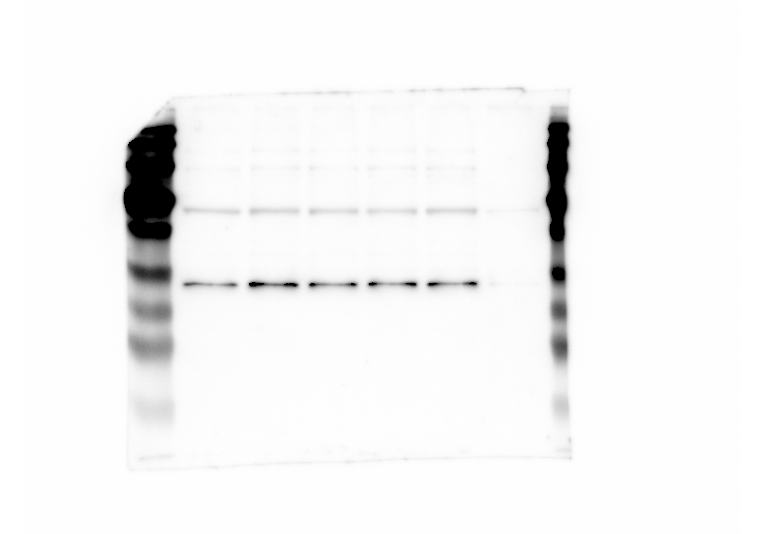

### Drp1 expression during preimplantation development.tif

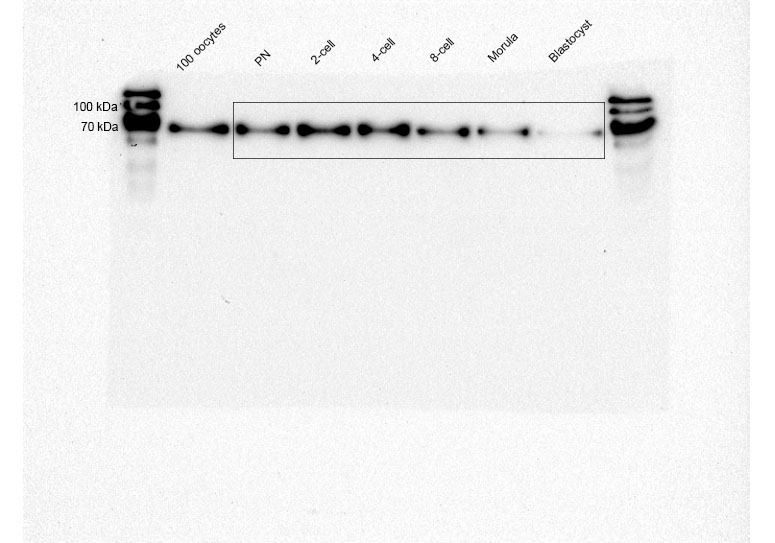

### Drp1 expression during preimplantation development_raw.jpg

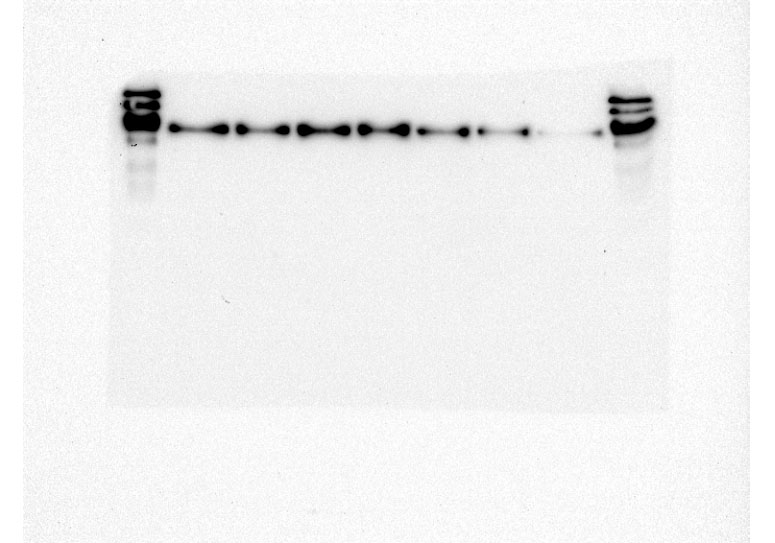

### Drp1 expression rescue raw.tif

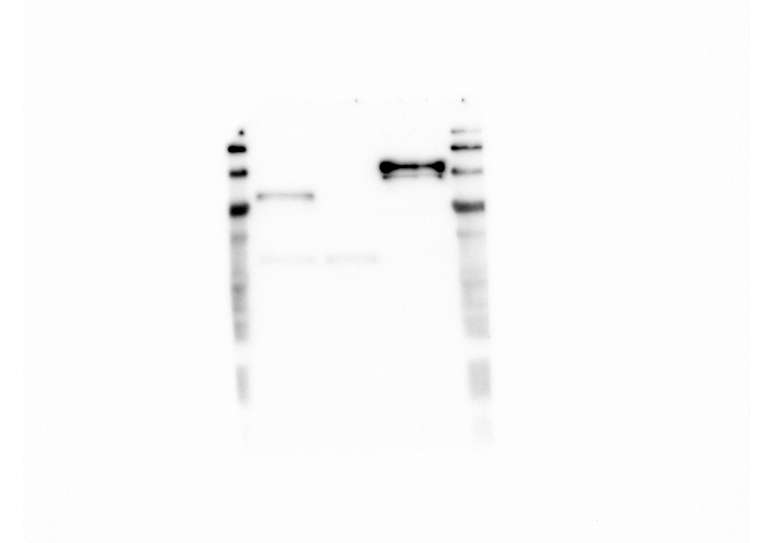

### Drp1 expression rescue.tif

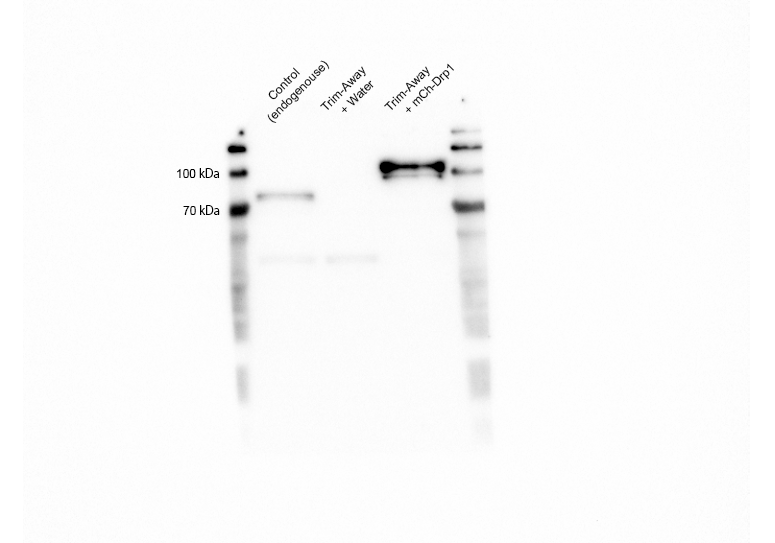

### Drp1 expression Trim Away raw.tif

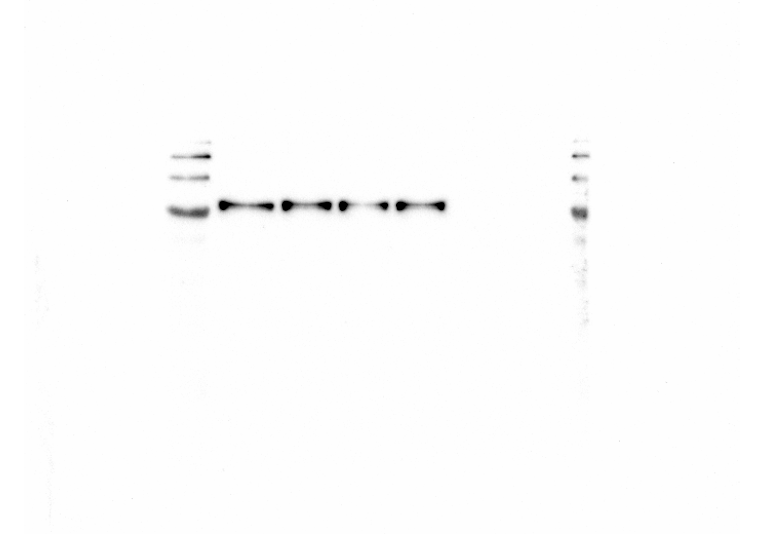

### Drp1 expression Trim Away.tif

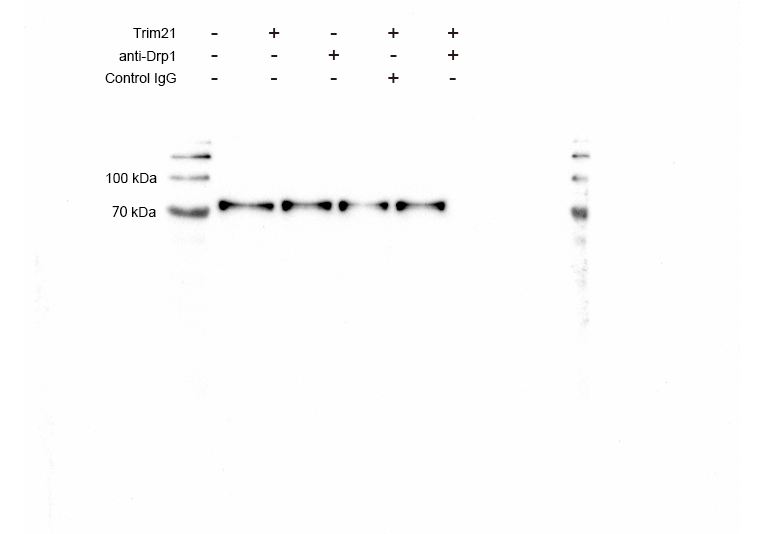

### Myo19 expression Trim Away a.tif

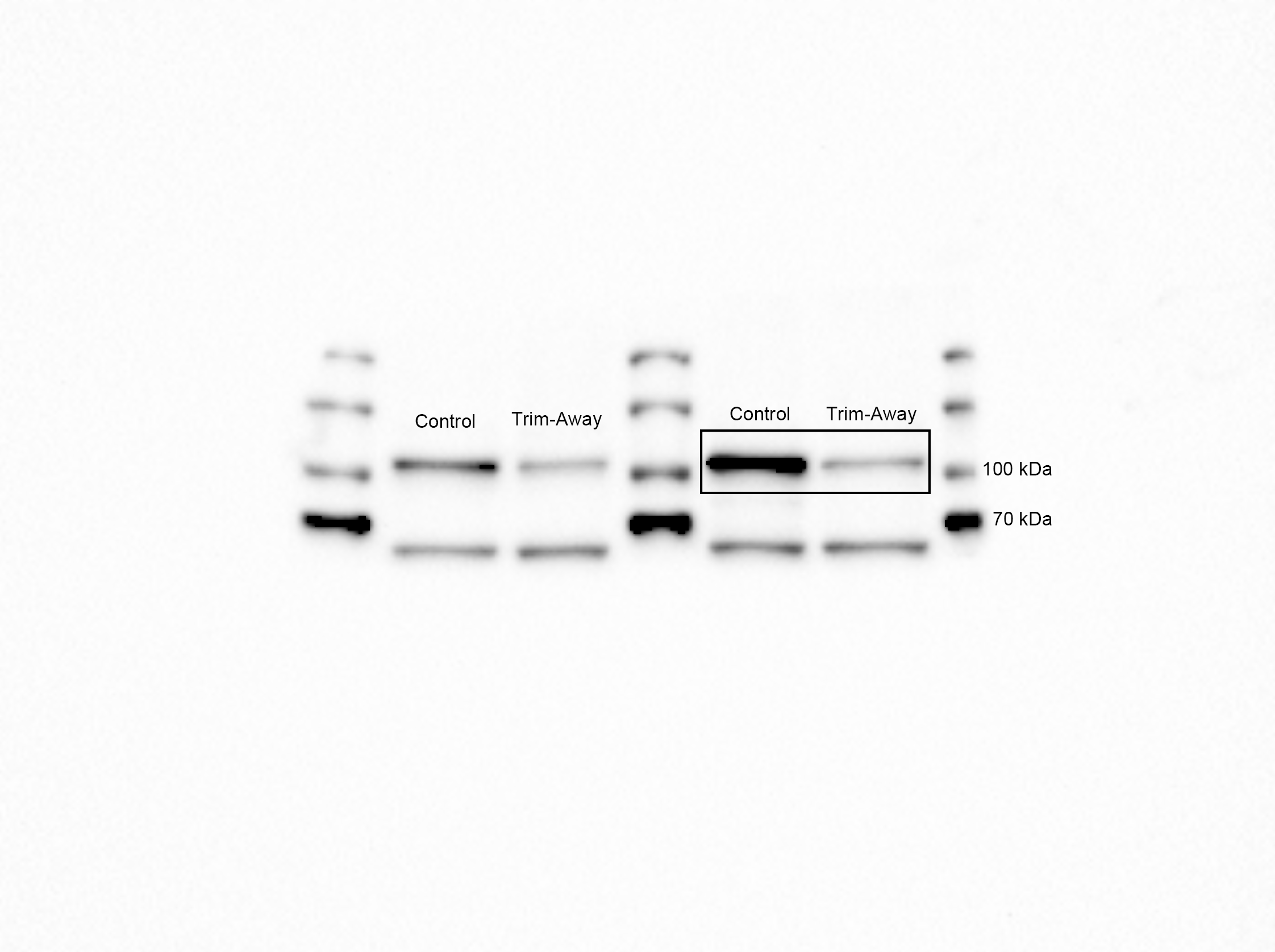

### Myo19 expression Trim Away raw.tif

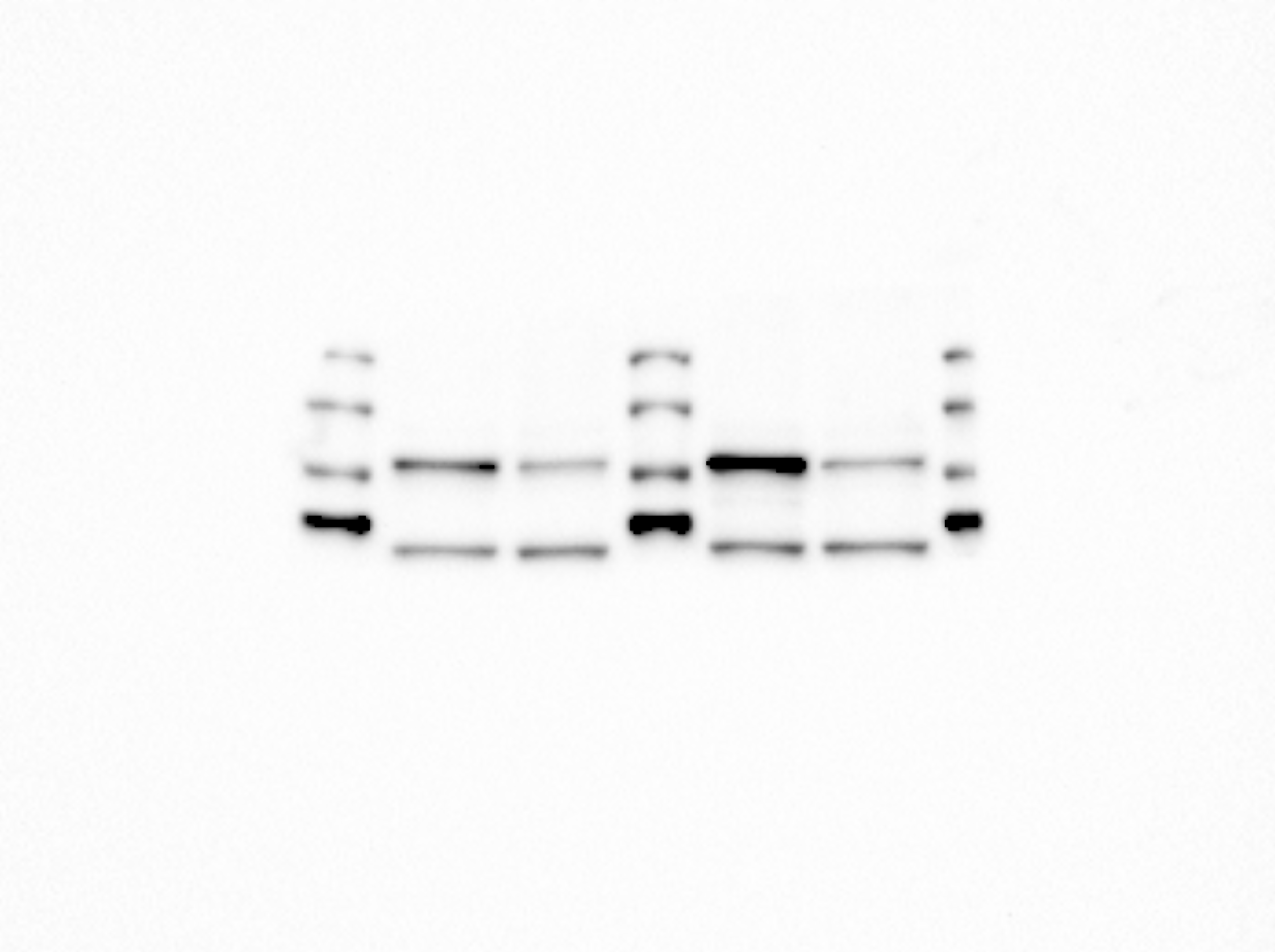
